# Mitochondrial haplotype and mito-nuclear matching drive somatic mutation and selection throughout aging

**DOI:** 10.1101/2023.03.06.531392

**Authors:** Isabel M. Serrano, Misa Hirose, Charles C. Valentine, Sharon Roesner, Elizabeth Schmidt, Gabriel Pratt, Lindsey Williams, Jesse Salk, Saleh Ibrahim, Peter H. Sudmant

## Abstract

Mitochondrial genomes co-evolve with the nuclear genome over evolutionary timescales and are shaped by selection in the female germline. Here, we investigate how mismatching between nuclear and mitochondrial ancestry impacts the somatic evolution of the mt-genome in different tissues throughout aging. We used ultra-sensitive Duplex Sequencing to profile ∼2.5 million mt-genomes across five mitochondrial haplotypes and three tissues in young and aged mice, cataloging ∼1.2 million mitochondrial somatic and ultra low frequency inherited mutations, of which 81,097 are unique. We identify haplotype-specific mutational patterns and several mutational hotspots, including at the Light Strand Origin of Replication, which consistently exhibits the highest mutation frequency. We show that rodents exhibit a distinct mitochondrial somatic mutational spectrum compared to primates with a surfeit of reactive oxygen species-associated G>T/C>A mutations, and that somatic mutations in protein coding genes exhibit signatures of negative selection. Lastly, we identify an extensive enrichment in somatic reversion mutations that “re-align” mito-nuclear ancestry within an organism’s lifespan. Together, our findings demonstrate that mitochondrial genomes are a dynamically evolving subcellular population shaped by somatic mutation and selection throughout organismal lifetimes.

## Introduction

The mitochondrial genome (mt-genome) encodes for 13 proteins that are vital for the electron transport chain. However, more than 1,000 nuclear encoded genes are necessary for mitochondrial assembly and function^1^. The required coordination between nuclear- and mitochondrial-encoded proteins drives the coevolution of these two genomes^2–4^. This concerted evolution has been observed in both laboratory crosses and natural hybrids across a variety of species, including: fruit flies^5,6^, marine copepods^7,8^, wasps^9–11^, yeast^12,13^, eastern yellow robins^14^, swordtail fish^15^, and teleost fish^16^. Hybrids often exhibit reduced fitness attributed to attenuated mtDNA copy number^17^, mtDNA gene expression^17,18^, and OXPHOS function^6–8,10^. In natural hybrids, the genetic ancestry at nuclear-encoded mitochondrial genes is highly differentiated between populations that have distinct mt-haplotypes^14^. In human admixed populations, mito-nuclear ancestral discordance has been shown to correlate with reductions in mtDNA copy number^19^. Additionally, *de novo* germline mitochondrial mutations in admixed human populations exhibit a bias towards concordance with nuclear ancestry^20^. In somatic tissues, the nuclear genome has been shown to influence the segregation of mtDNA haplotypes in a tissue-specific manner^21–23^. Together, these studies highlight the functional importance of mito-nuclear ancestral concordance. However, the impact of these genomic interactions on somatic mutation and selection has not been studied.

As healthy cells age, they accumulate nuclear and mtDNA damage as a result of environmental exposures and cellular processes^24^. mtDNA has a 10- to 100-fold higher *de novo* germline mutation rate than nuclear DNA^20,25,26^ due to its lack of protective histones, a higher replication rate, and less effective DNA damage repair mechanisms^27^. Individual cells can contain hundreds to thousands of mt-genomes^28,29^, presenting a dynamic subcellular population that is highly susceptible to mutation. While the relationship between mutations in the mt-genome and aging remains unclear^30,31^, increased mtDNA damage has been associated with many aging phenotypes and several age-related diseases^30,32^. Yet, profiling low frequency mutations in this population of mt-genomes has historically been challenging with most technologies limited to variants segregating at a frequency >1%^26,33^. More recently, several studies have employed Duplex Sequencing^33^ to capture mutations with an error rate of <1×10^-7^. These works have examined several different species and confirmed a robust age-associated increase in mtDNA somatic mutation frequency^34–37;^ identified that mitochondrial somatic mutations stem from the replication process either via DNA polymerase gamma misincorporations or spontaneous DNA deamination of cytosine and adenine ^34–37;^ and observed tissue-specific somatic mutation rates^36,37^.

Here, we investigate the impact of mitochondrial ancestry (mt-ancestry) and tissue-type on the somatic evolution of the mt-genome through age. To explore how mitochondrial haplotype and mito-nuclear genomic interactions shape the mt-genome mutational landscape, we utilize a panel of mouse strains that have identical nuclear genomes but differ in their mitochondrial haplotypes (conplastic strains). We sample the brain, heart, and liver of these conplastic strains and the wildtype (C57BL/6J) in young and aged individuals. These tissues are physiologically distinct yet some of the most metabolically active, allowing us to study how evolutionary processes unfold in different molecular contexts. We use Duplex Sequencing to profile mt-genomes at an unprecedented level of depth and accuracy, allowing us to identify mutational hotspots and characterize the mutational spectrum. We use these results to discern signatures of selection acting to shape the mt-genome through age and confirm the existence of somatic mutations that work to realign mito-nuclear ancestry at short, within-lifespan, evolutionary timescales. Together, these findings characterize somatic evolution in the context of an organelle implicated in aging and age-related phenotypes.

## Results

### Ultra-sensitive duplex sequencing of 2.5 million mitochondrial genomes allows profiling of mutations to a frequency of 4×10^-6^

To investigate how mitochondrial haplotype and mito-nuclear concordance influence the distribution of mt-genome somatic mutations, we employed a panel of conplastic mouse strains. These strains are inter-population hybrids developed by crossing common laboratory strains with the C57BL/6J (B6) mouse line^38^. Each conplastic mouse strain carries a unique mitochondrial haplotype on a C57BL/6J nuclear genomic background (**Fig. 1 A**). This fixed nuclear background enables us to attribute differences in somatic mutation to changes in mitochondrial haplotype. Alongside a wildtype B6 mouse, we used four conplastic mouse strains that exhibit changes in metabolic content and processes, or altered aging phenotypes **(Table S1)**. Three of these conplastic mouse strains differ from the C57BL/6J mitochondrial haplotype by just 1-2 nonsynonymous variants, while a single strain (NZB) contains 91 variants distributed across the mt-genome **(Fig. 1B, Table S1)**. We sampled brain, liver, and heart tissues from each of these mouse strains in young (2-4 months old (mo.)) and aged (15-22 mo.) individuals to examine how aging and metabolic demands shape the mutational spectrum in distinct physiological contexts. In total, three tissues were sampled from five mouse strains at two different ages over 3-4 replicates per condition, resulting in n=115 samples.

**Figure 1.**
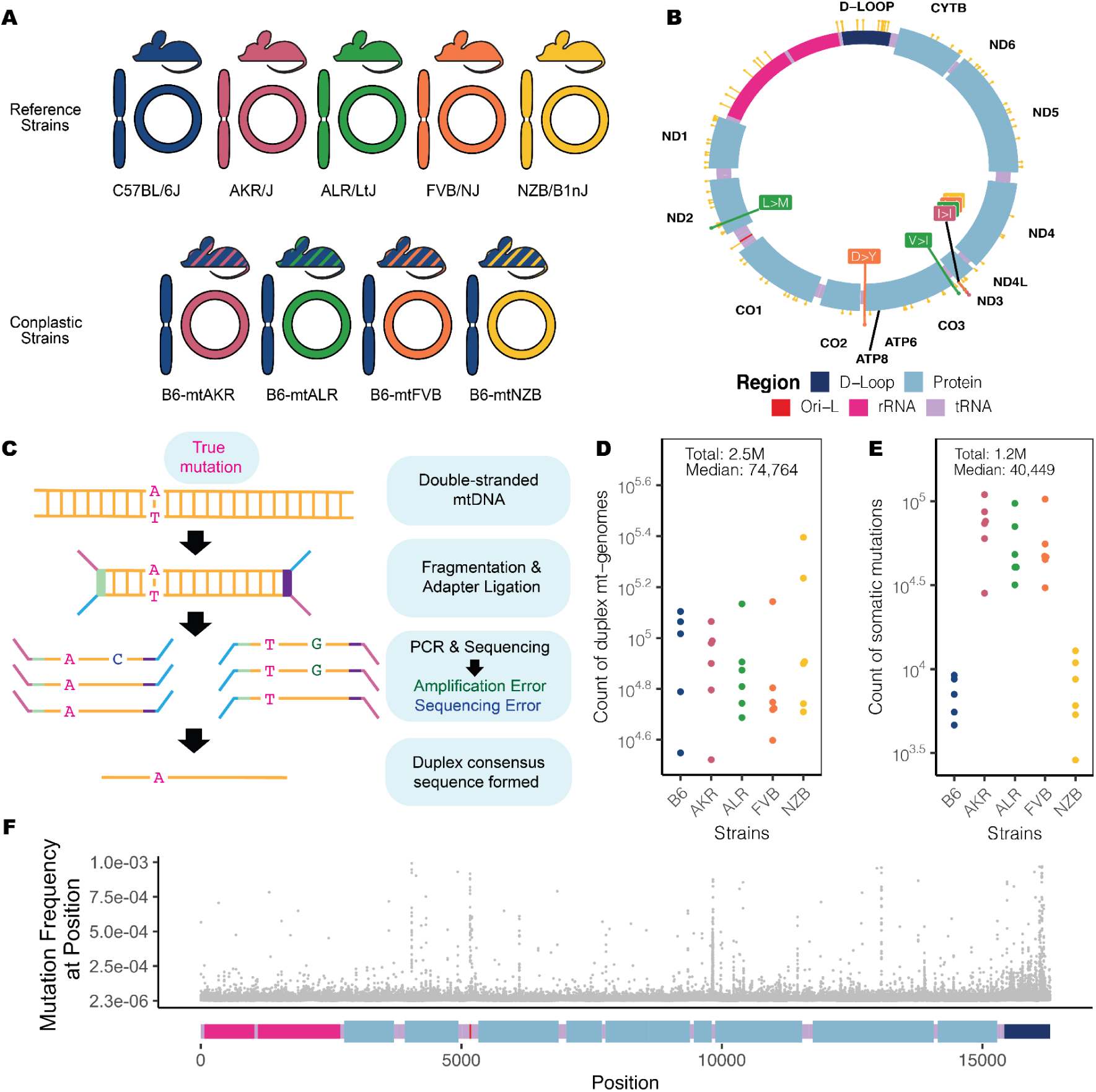
Overview of experimental design. (A) Maternal donors were selected from common inbred strains (CISs) containing nonsynonymous variants with functional effects. Females from these CISs (pink, green, and orange) were backcrossed with a C57BL/6J mouse (blue). This process was repeated with a wild-inbred strain (yellow). The result of these crossings are conplastic mouse strains (denoted by the striped mice). These conplastic strains have identical B6 nuclear (linear) genetic backgrounds but differ by variants along their mt-genomes (circular). (B) B6-mtAKR (pink), B6-mtALR (green), and B6-mtFVB (orange) differ from wildtype by 1-3 missense variants. B6-mtNZB (yellow) contains 91 variants distributed across the mt-genome (14 missense and 56 synonymous mutations). The synonymous mutation in *mt-ND3* is shared across conplastic strains. For a more detailed account of these variants and their associated phenotypes reference **(Table S1)**. (C) Ultra-sensitive Duplex Sequencing (DupSeq) was used to profile mt-genomes from conplastic mice. DupSeq has an unprecedented error rate of <10^-7^. Each double-stranded mtDNA fragment is distinctly tagged with a unique molecular barcode, allowing for the computational construction of a duplex consensus sequence. As a result, both PCR and sequencing errors are filtered from the data. (D) The count of duplex mt-genomes sequenced was calculated for each experimental condition (n = 29 conditions; n = 4 mice for every condition with 3 mice for B6-Young-Heart). The duplex read depth at each position was aggregated across samples in a condition to quantify the duplex depth per condition. The average duplex read depth across the mt-genome was then calculated. An estimated 2.5 million mt-genomes were duplex sequenced with a median of 74,764 mt-genomes duplex sequenced per condition. (E) Approximately 1.2 million somatic variants were identified with a median of 40,449 variants per condition. The count of somatic variants was aggregated across samples in a condition. Mutations present at conplastic haplotype sites were filtered from the analysis. (F) The mutation frequency for each position is mapped along a linear representation of the mt-genome. Each point denotes the mutation frequency for an experimental condition at the given position in the mt-genome. Legends for the mt-genome map are consistent with those in **(Fig. 1B)**. Positions with a mutation frequency greater than 1×10^-3^ were excluded from this analysis.

In order to capture low frequency variants and accurately portray the somatic mutational landscape, we used ultra-sensitive Duplex Sequencing to profile mt-genomes across different experimental conditions^33^. This approach works by tagging double stranded mtDNA sequences with molecular identifiers and computationally constructing a duplex consensus sequence **(Fig. 1C)** resulting in error rates of ∼2 × 10^-8^. Importantly, duplex sequencing data can contain an artificial enrichment of G>T/C>T and G>C mutations resulting from DNA damage during sequencing preparation steps^39^. To exclude potential erroneous calls, we trimmed 10 bp from our duplex reads **(Supplementary Note 1, Fig. S1).** Additionally, we analyzed the mutational spectra for atypical imbalances of complementary mutation types and compared our mutation type proportions to a previously published study^37^ **(Fig. S2, Table S2, Fig. S3, Table S3)**. Using this approach, we profiled ∼2.5 million duplex mt-genomes, with a median of 74,764 duplex mt-genomes sequenced per condition (*strain x tissue x age*) **(Fig. 1D)**. This resource allows for the detection of somatic mutations at a frequency of 4×10^-6^.

Duplex reads from each sample were mapped to the mouse mitochondrial genome (mm10) and variants were called using a Duplex Sequencing processing pipeline (see Methods). Mutations overlapping conplastic haplotype sites were filtered out. In total we identified 1,171,918 somatic variants, with a median of 40,449 somatic mutations per condition **(Fig. 1E).** From these ∼1.2 M mutations, 81,097 mutations are unique events. These variants were distributed across the entirety of the mt-genome **(Fig. 1F)**.

### Haplotype- and tissue-specific mitochondrial mutation rates and hotspots

Somatic mutations accumulate with age across both nuclear and mitochondrial genomes^24^. We observed this trend consistently across tissues and mt-haplotypes with the mutation frequency on average ∼2-fold higher in aged mice compared to young mice **(Fig. 2A)**. This trend was most pronounced in the liver (2.5-fold higher) and smallest in the heart (1.6-fold higher). Additionally, the heart sustained the lowest mutation frequency on average **(Fig. 2A)**, despite its high metabolic demands as demonstrated by its higher mitochondrial copy number **(Fig. S4)**.

**Figure 2.**
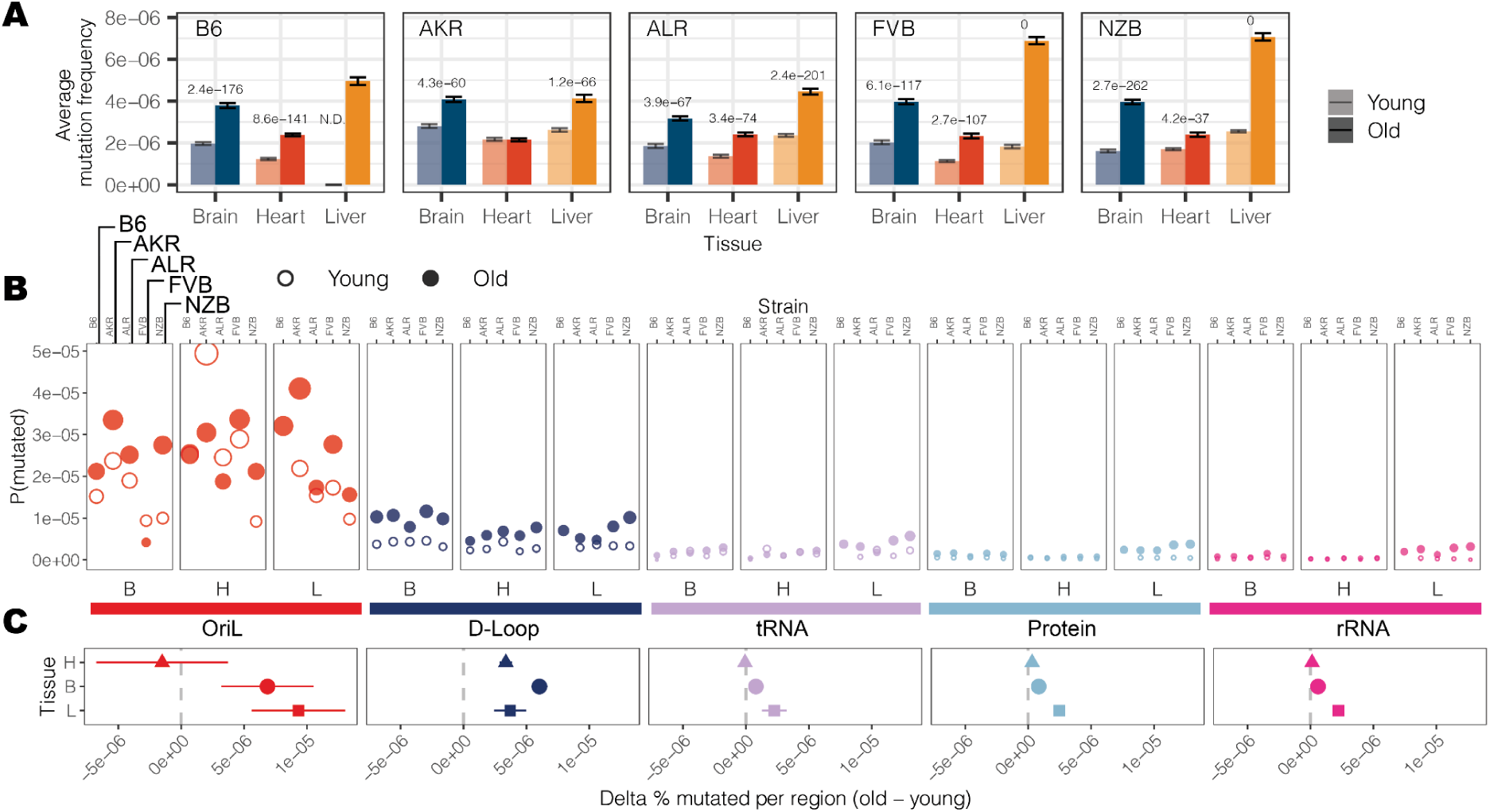
Region-specific changes in somatic mutation frequency with age. (A) The average mutation frequency for each experimental condition. The mutation frequency for each position along the mt-genome was calculated by dividing the total count of alternative alleles by the duplex read depth at the position. Error bars denote the 95% Poisson confidence intervals. P-values indicate experimental conditions with a significant age-associated increase in somatic mutation frequency (log-linear regression and adjusted p-values using a Bonferroni multiple hypothesis correction). (B) The mt-genome was categorized into regions: OriL (red), D-Loop (dark blue), tRNAs (purple), protein coding (light blue), and rRNAs (magenta). For each region, the probability of mutation was calculated as the total count of mutations normalized by the region length in bp multiplied by the average duplex read depth across the region. Fill indicates age group: young (hollow circles) and aged (filled circles). (C) The difference in percent bp mutated between young and aged mice. The shape denotes the average difference in percent bp mutated with age across conditions. Error bars showcase the standard error of the mean. Positions that exceeded a mutation frequency of 1×10^-3^ were excluded from these analyses. For Fig. 2B and 2C, frequencies were normalized for sequencing depth across conditions at each position.

Comparing age-associated mutation rates across different strains we find AKR, ALR, and NZB all exhibit strain-specific mutation rates **(Table S4)**, p-value < 0.01: log-link regression). The FVB strain, which only differs from B6 at two sites **(Fig. 1B)** showed no evidence of strain-specific mutation rates compared to wildtype. AKR and ALR strains had lower mutation rates across all tissues while NZB exhibited strong increases in the brain and decreases in the heart.

We next examined the variation in mutation rates and frequencies across different regions of the mitochondrial genome **(Fig. 2B, C)**. The density of mutations was lowest in functional coding regions (protein coding, rRNA, and tRNA segments) while the *D-loop* exhibited a significantly higher mutation frequency in both young (6.4-fold, p-value = 3.4 × 10^-9^; two-tailed t-test) and aged (4.5-fold, p-value = 8.6 × 10^-8^; two-tailed t-test) mice. However, we further found that the light strand origin of replication (*OriL*) had an even higher average mutation frequency, 40- to 22-fold in excess of the functional coding regions in young and aged mice, respectively **(Fig. 2B)**. This *OriL* hotspot of mutation was most pronounced in aged wildtype B6 mice. The *OriL* was recently noted as a mutational hotspot in macaque liver, but not in oocytes or muscle in that species^36^. In contrast, we find the *OriL* to exhibit elevated mutation frequencies across the brain, heart, and liver.

Our initial inspection of the overall distribution of mutations across the mt-genome highlighted several distinct clusters appearing at finer-scale resolution than our simple functional classification groups **(Fig. 1F).** To resolve these putative mutational hotspots we quantified the average mutation frequency in 150 base-pair (bp) sliding windows over the mt-genome independently across tissues and mt-haplotypes **(Fig. 3A)**. In addition to the *D-loop*, this analysis identified mutational hotspots in *OriL, mt-ND2*, and *mt-tRNA^Arg^*.

**Figure 3.**
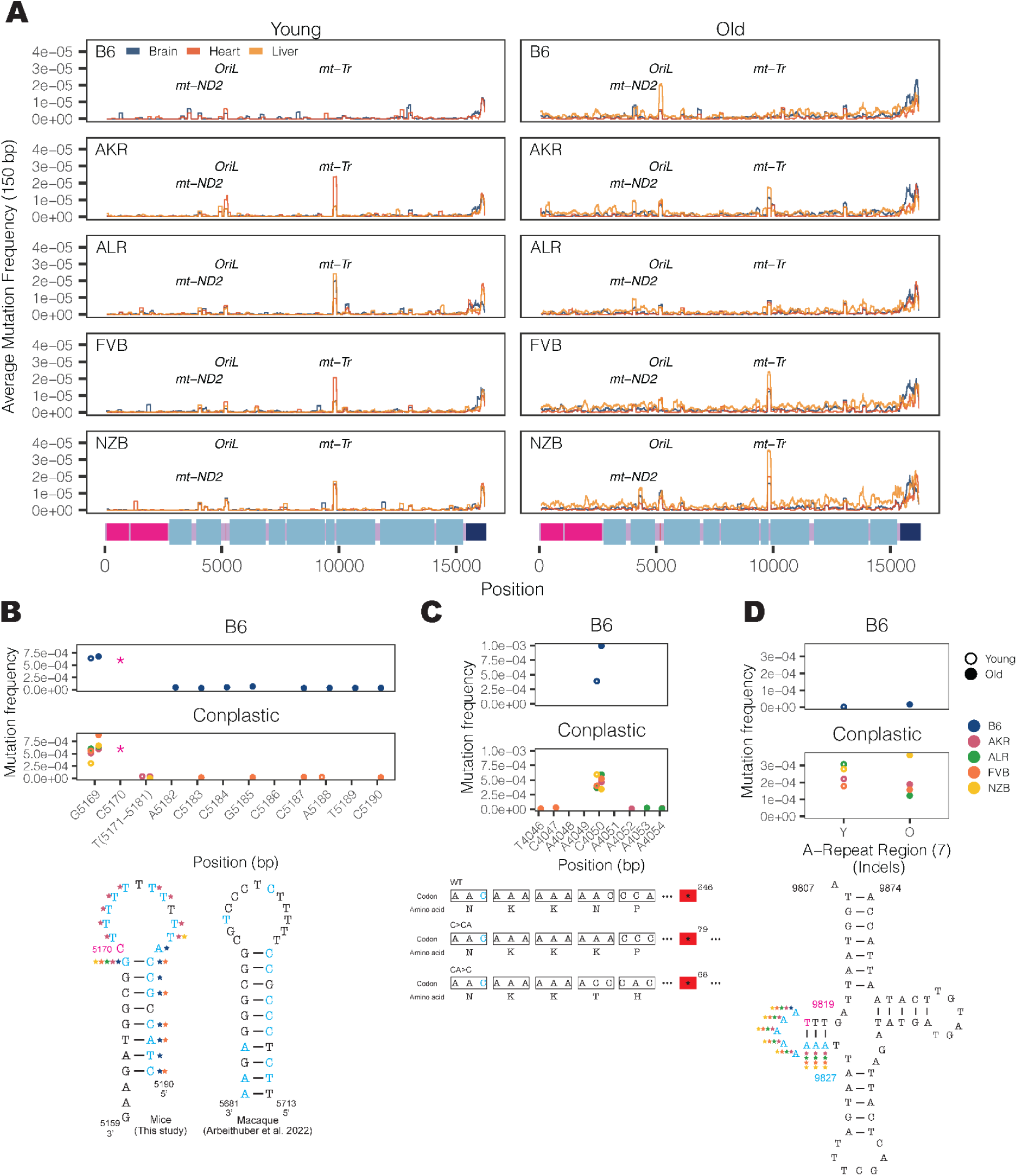
Haplotype-specific peaks of mutation along the mt-genome. (A) The average mutation frequency was calculated in 150-bp sliding windows for each strain and tissue independently. Each line represents a tissue: brain (blue), heart (red), and liver (orange). Regions with shared mutation frequency peaks in at least three mouse strains are labeled. (B) The high frequency region in OriL is highlighted. For positions 5171-5181 the mutation frequency for the T-repeat region is calculated as the sum of mutations across this region divided by the sum of the duplex depth across positions. Each color denotes a different strain: B6 (blue), AKR (pink), ALR (green), FVB (orange), NZB (yellow). The average frequency in young (left, hollow points) and aged (right, filled points) mice are compared. The schematic compares the *OriL* structure in mice to that of macaques^36^. Positions with a mutation frequency > 1×10^-3^ are in magenta, while positions that undergo mutations are denoted in light blue. In the macaque *OriL* diagram, variant hotspots are denoted in light blue. Stars represent strains that have mutations present at a given position. (C) The high frequency region in *mt-ND2* is highlighted. Color, shape, and calculation of the average mutation frequency are similar to those in Fig. 3B. A schematic demonstrating the sequence and codon changes that result from the indels at position 4050: premature stop codons at codon 79 and 68 for C>CA and CA>C, respectively. The superscript denotes the position of the premature stop codon in the amino acid sequence. (D) The mutation frequency for *mt-tRNA^Arg^* from bp positions 9820-9827, which is an A-repeat region. The mutation frequency for the A-repeat region is calculated as the sum of mutations across this region divided by the sum of the duplex depth across positions. The diagram of *mt-tRNA^Arg^* highlights in magenta the location of a fixed variant in NZB which is a high heteroplasmic variant in the other mouse strains. Positions that have mutations in this region are in light blue. Positions that exceeded a mutation frequency of 1×10^-3^ were excluded from these analyses. Frequencies were normalized for sequencing depth across conditions at each position.

The *OriL* forms a stem loop structure that is conserved across species^40^ however differs markedly in its sequence. We identified mutations throughout the loop and 3’ end of the stem with the highest mutation frequencies corresponding to changes in the size of the stem loop (**Fig. 3B**). Repeated mutations of the 3′ end of the stem loop structure mirror those observed in macaque liver in a recent study^36^, though the sequence of the stem differs between these two species. Thus, the conserved structure of the *OriL* appears to drive convergent mutational phenotypes between species.

The *mt-ND2* mutation hotspot consists of a frameshift-inducing A insertion or deletion that introduces a premature stop codon **(Fig. 3C)**. This stop codon reduces the length of the final protein product by more than 250 amino acids, likely severely impacting its function. This mutation increases in frequency with age across all strains except NZB. The final mutation hotspot we identified is localized to an 8 nucleotide stretch of *mt-tRNA^Arg^*(**Fig. 3D**). These nucleotides correspond to the 5′ D-arm stem-loop of the tRNA and constitute primarily A insertions 2-4 nucleotides long. Computational tRNA structure predictions show that these mutations increase the size of the loop **(Fig. S5, Fig. S6)**. This mutational hotspot exhibits strong strain-specificity with B6 exhibiting few somatic mutations in this A-repeat stretch. The D-arm plays a critical role in creating the tertiary structure of tRNAs ^41,42^ potentially contributing to its constraint in the wildtype. Together, these results highlight the emergence of distinct mutational hotspots in mitochondrial genomes occurring in age- and strain-specific contexts, and implicating both DNA and RNA secondary structures.

### DNA replication error and deletion events are distinguishing features of aged mitochondrial genomes

Somatic mutations are caused by various molecular processes that lead to DNA damage, each of which exhibit a distinct mutational signature^43^. To identify sources of mitochondrial somatic mutation, we categorized mutations into single nucleotide variants (SNVs), deletions (DEL) and insertions (INS) **(Fig. 4A)**. SNVs were further classified into the six possible substitution classes **(Fig. 4B)**. Overall, SNVs were the predominant somatic mutation type, with a 5-fold higher average mutation frequency than deletions and insertions in both young and aged individuals **(Fig. S7)**. The most abundant mutation type observed was G>A/C>T, which is indicative of replication error or cytosine deamination to uracil^44,45^ **(Fig. 4B)**. The reactive oxygen species (ROS) damage signature of G>T/C>A^46^ was the second most predominant mutation type. All somatic mutation types were used to identify two dominant mutational signatures using a multinomial bayesian inference model^47^ **(Fig. 4C)**. These signatures explained the bulk of the variation across samples **(Fig. S8)**.

**Figure 4.**
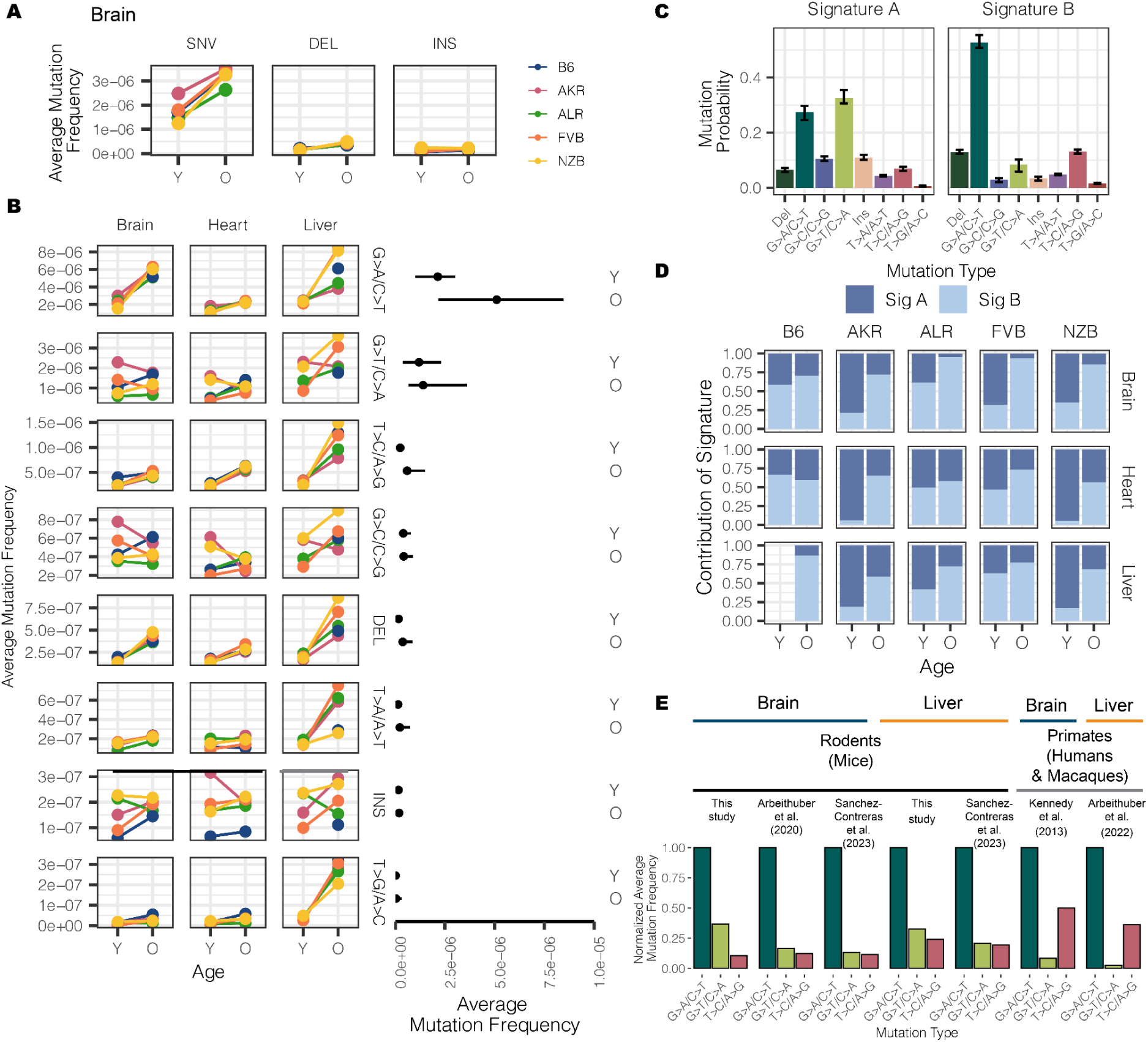
Characterizing the mt-genome mutational landscape. (A) The average mutation frequency is compared between mt-haplotypes for different classes of mutations: single nucleotide variants (SNVs), deletions, and insertions. The average mutation frequency was calculated as the total mutation count in a class divided by the total duplex bp depth. Results for the brain are featured, which showcase trends observed in the heart and liver, as well **(reference Fig S7)**. (B) SNVs were further classified into point mutation types. The average mutation frequency for each mutation type was compared across mouse strains. The average frequency for point mutations was calculated as the total count of mutations of the given type divided by the duplex bp depth for the reference nucleotide. The average mutation frequency for deletions and insertions was calculated as the total mutation count in these classes divided by the total duplex bp depth. Mutation types are ordered by descending mutation frequencies. The range in frequency across experimental conditions for each mutation type is illustrated by the line segments and compared across age groups. Each point in the segment denotes the median frequency across conditions. (C) Mutational signatures were extracted from mutation type counts for each experimental condition using sigfit. Two mutational signatures were identified. The probability that a mutation type contributed to a signature is showcased with error bars denoting the 95% confidence interval of each probability. (D) The presence of the mutational signatures was estimated for each condition (Signature A dark blue, Signature B light blue). The contribution of each mutational signature is compared across age. (E) The frequency of the three most abundant mutation types were compared across mice, macaques, and humans for the brain and liver. Mutation frequencies were normalized by the frequency of the G>A/C>T mutation in their respective study. All studies used Duplex Sequencing to profile mutations in the mt-genome. For the mutation frequencies in our study, we only show the mutation frequencies for B6. Refer to **Fig. S9A** for frequencies across all strains, tissues, and young mice. For all analyses, mutation counts and duplex bp depth were aggregated across samples in an experimental condition. Positions with mutation frequencies greater than 1×10^-3^ were omitted from these analyses.

The accumulation of somatic mutations with age is a well known phenomenon that has been hypothesized to play key roles in the etiology of lifespan^48^. We observed that 4 out of the 8 mutation types significantly increased in frequency with age (adjusted p-value < 0.01; Fisher’s Exact Test with a Benjamini-Hochberg correction) **(Fig. 4B, Table S5)** across all tissues, with T>A/A>T mutations additionally exhibiting age-associated accumulation in the liver but not in brain or heart. These age-associated signatures compose *Mutational Signature B* **(Fig. 4C)**, which overall distinguishes aged from young samples **(Fig. 4D)**. The consistency of these signatures across both mutations and mt-haplotypes suggest that these mutations occur in a ’clock-like’ fashion in mitochondria over lifespan.

In contrast, insertions, G>C/C>G, and G>T/C>A mutations did not exhibit consistent age-associated patterns. These mutations are represented in *Mutational Signature A* **(Fig. 4C)**. Of particular note, the G>T/C>A and G>C/C>G mutation patterns are associated with ROS damage. An increase in ROS damage has been hypothesized to play an important role in aging. Nonetheless, we do not find evidence of an increase in ROS-associated damage with age. Previous works in the human brain^34,49^, macaque tissues^36^, and various mouse tissues^35,37^ have also observed a lack of ROS-associated damage with age.

### Evidence of distinct somatic mitochondrial mutational signatures across species

While the overwhelming majority of research into somatic mutation rates and profiles has focused on humans, comparative analyses can provide insight into the evolutionary processes that have shaped mutation. To determine if mitochondrial somatic mutation exhibited species-specific patterns we compared several recent studies that Duplex Sequenced mt-genomes in multiple mouse, human, and macaque tissues^34–37^. In our dataset, we found the relative magnitude of mutation frequencies to be consistent across tissues and mt-haplotypes with the G>A/C>T and G>T/C>A mutations being the most abundant **(Fig. 4B, Fig. S9A).** This signature is consistent with other duplex sequencing-based analyses of mitochondrial mutations in young and aged mouse brain, muscle, kidney, liver, eye, heart, and oocytes **(Fig. 4E, Fig. S9B, Fig. S9C)**^35,37^. However, profiling of mitochondrial mutation signatures in several young and aged human brains noted transitions, G>A/C>T and T>C/A>G, to be the most abundant mutation signatures^34^ **(Fig. 4E)**. We find that this pattern is also recapitulated in a recent dataset of mitochondrial somatic mutation in macaque muscle, heart, liver and oocytes **(Fig. 4E, Fig. S9D)**^36^. These differences in mutational signatures were observed between primate and rodent species from duplex sequencing data generated independently by two research groups, emphasizing the reproducibility of this phenomenon. Together, these results suggest that rodent and primate mt-genomes are subject to distinct mutational processes, potentially as a cause or effect of physiological differences between these lineages.

### Purifying selection persists among artificial signals of positive selection

While only thirteen proteins are encoded on the mt-genome, these products are vital components of the electron transport chain. Given their importance, we investigated whether selection was acting to shape the mutational landscape of the mt-genome.

We calculated the 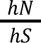 statistic^50^, which is akin to the 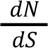 statistic, for every gene across our 29 experimental conditions. For each condition, we also simulated the expected distribution of 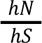 ratios using the observed mutation counts and mutational spectra **(Fig. S10)**. *Prima facie*, the mt-genome appeared to be shaped predominantly by positive selection (**Fig 5A**). However, differences in the mutation spectrum between mutational hotspots, including the D-loop, could bias the simulated null distribution of mutations. Indeed, mutation spectra were significantly different between D-loop and non-D-loop mutations, as well as among mutations at different frequencies (**Fig 5B**). Furthermore, quantifying the allele frequency spectrum of missense and synonymous mutations revealed that missense substitutions were strongly enriched at low frequencies compared to synonymous substitutions, indicating negative selection dominating at higher frequencies (**Fig 5C**). We thus performed our 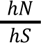 analyses independently in different frequency bins (**Fig 5A**) revealing primarily signatures of negative selection. From the twelve possible cases of selection, 75% were in conplastic mice, and both cases of positive selection were in NZB mice. Genes with selective signatures include *COXI* (3), *COXIII* (1), *Cytb* (1), *ND2* (1), and *ND5* (6), which compose complexes I, III, and IV of the electron transport chain.

**Figure 5.**
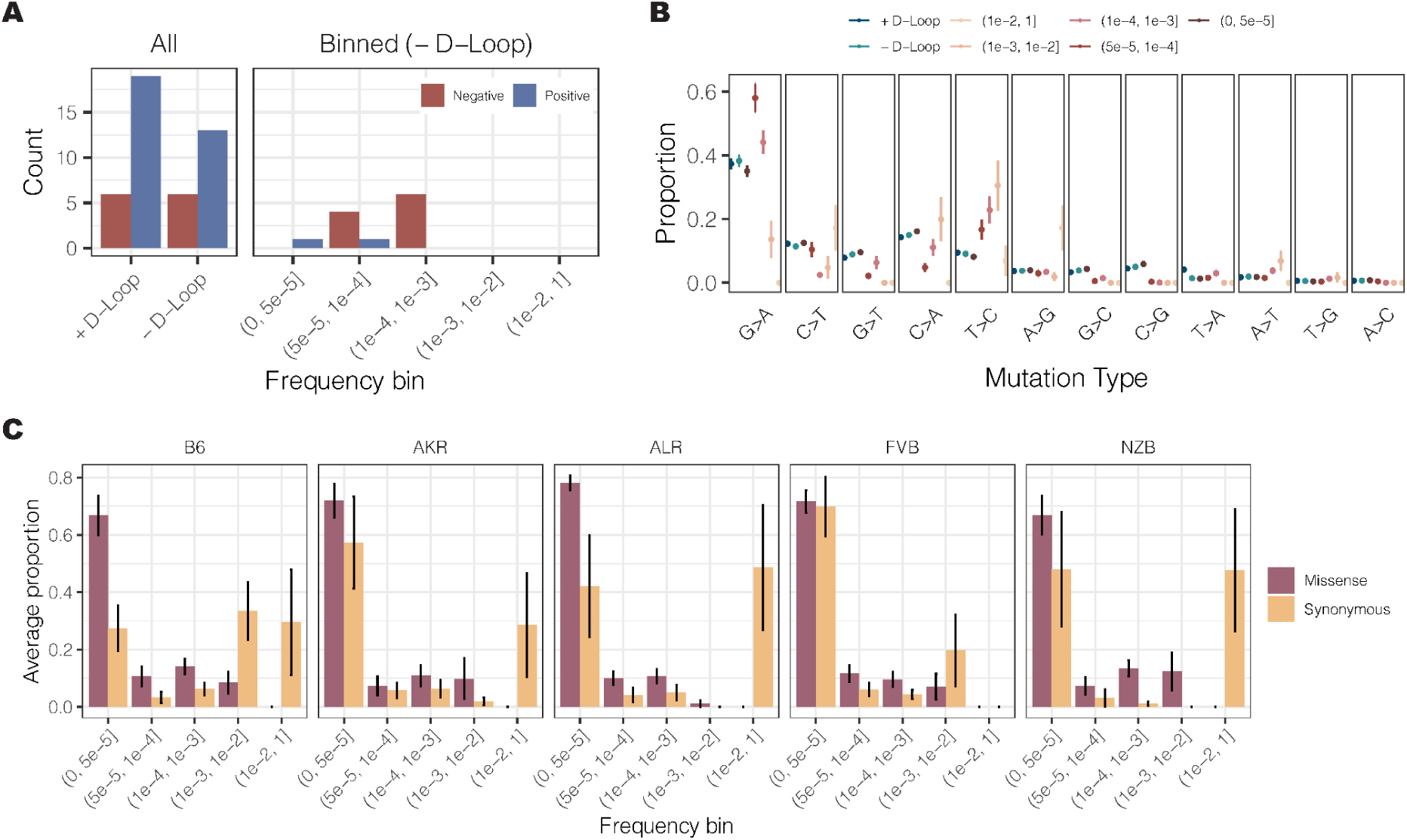
Aggregating mutation type proportions across frequency bins induces a false signal of positive selection in the mt-genome. (A) The count of genes under positive (blue) and negative (red) selection. All constitutes analyses where the observed mutations and mutation types across frequencies were used to simulate 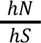 statistics. Mutations with a frequency greater than 1 × 10^-3^ were excluded from aggregated selection analyses. The analysis was also conducted for mutations binned according to their frequencies. For the binned analysis, mutations residing in the D-Loop were excluded. (B) The proportion of each mutation type. Proportions are calculated as the count of mutations of each type divided by the total count of mutations in a given bin. The average mutation type proportion across experimental conditions is shown with error bars denoting the standard error of the mean. In blue are proportions for the aggregated mutation counts with (dark blue) and without (light blue) the D-Loop. Mutations are ordered in descending order of average mutation frequency. (C) The frequency spectra for missense (purple) and synonymous (orange) mutations. The proportion denotes the count of missense or synonymous mutations in a frequency bin relative to the total count of missense or synonymous mutations. The average proportion across tissues and age in a strain is shown with error bars denoting the standard error of the mean.

Intriguingly, signatures of negative selection in *ND5* were observed across all five strains in our experiment. Taken together, these results indicate that negative selection dominates the distribution of mutations in mitochondrial genomes, though this signature is predominantly found at intermediate frequencies.

### Somatic reversion mutations are highly enriched and increase in frequency with age

Mismatching of mitochondrial and nuclear haplotypes, such as in hybrid populations, has been associated with reductions in fitness^5–13,15^. The conplastic strains we employ are hybrids with mismatched nuclear and mt-genome ancestries. Since hybrids often demonstrate reduced fitness as a result of mito-nuclear discordance, we reasoned that at sites where the conplastic mt-genome differed from the B6 mt-genome (haplotype sites), there may be a preference for “reversion” mutations to the B6 allele **(Fig. 6A)**.

**Figure 6.**
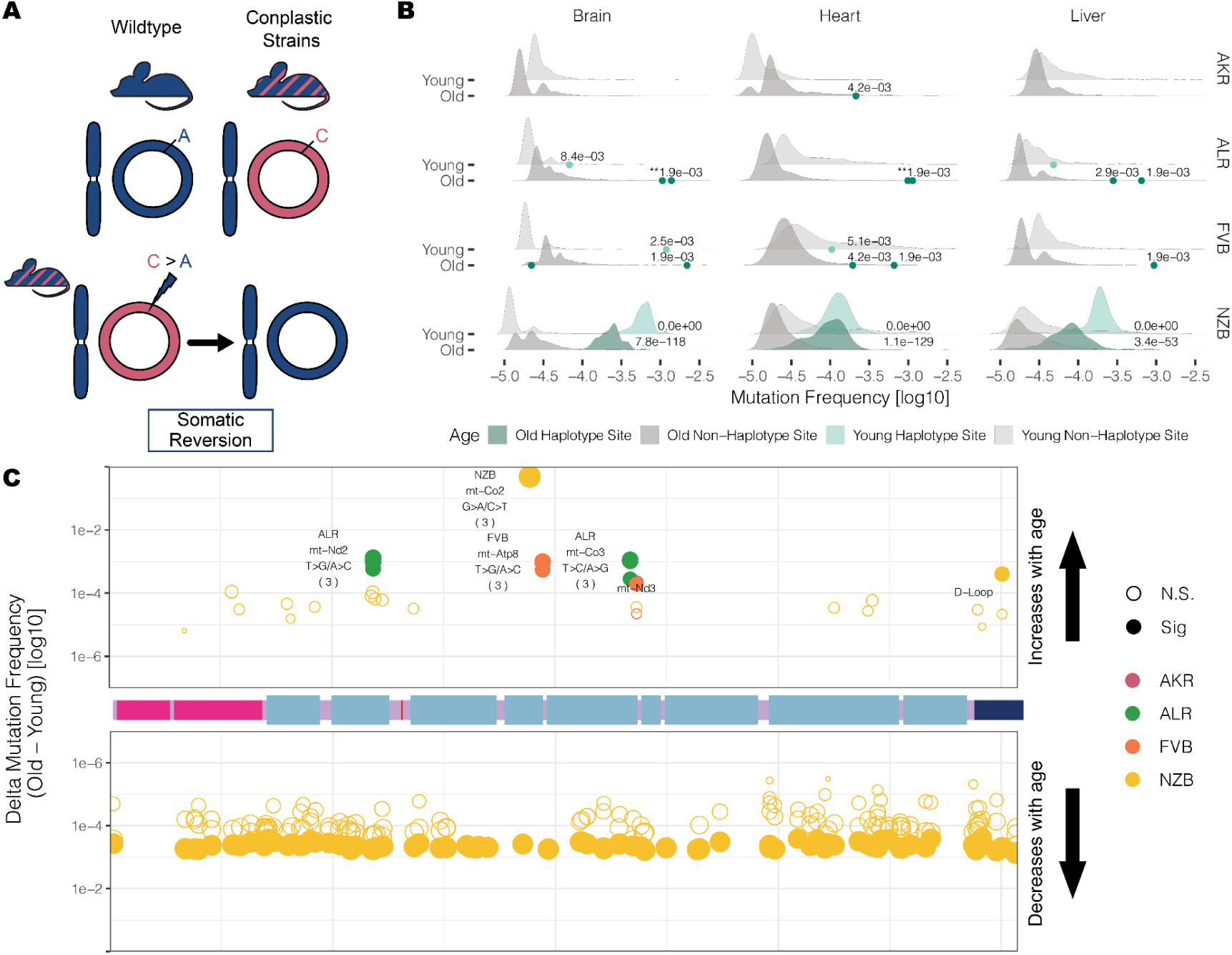
Somatic reversion mutations segregate in the mt-genome population. (A) A schematic that explains somatic reversion mutations. The wildtype strain (B6) has matching nuclear and mt-genome ancestries (B6 ancestry denoted in blue). Conplastic strains are hybrids with mismatching nuclear and mt-genome ancestry. Haplotype sites are positions in the mt-genome where the conplastic mt-genome differs from the B6 mt-genome. Somatic reversion mutations refer to the reintroduction of B6 ancestral alleles at haplotype sites. (B) The mutation frequency at non-haplotype sites (gray distributions) is compared to the mutation frequency at haplotype sites (green distributions or points when less than three haplotype sites exist). All mutations were included in the distribution of non-haplotype site mutation frequencies, including positions with a mutation frequency greater than 1×10^-3^. Haplotype site mutation frequencies were corrected for potential NUMT contamination. Mutation frequencies were normalized for sequencing depth between young and aged experimental conditions. Denoted are the adjusted empirical p-value. Asterisks denote the number of haplotype sites with the same p-value. (C) A map of the somatic reversion mutations along a linear mt-genome. Each point denotes the change in frequency with age (delta) for a somatic reversion mutation. The size of the point indicates the magnitude of delta and color represents the conplastic strain the somatic reversion occurs in. Fill denotes significance (adjusted empirical p-value < 0.01).

We hypothesized that if somatic selection were to favor the B6 allele, then reversions would occur at a higher rate than background mutations **(Table S6)**. Three of the strains have several fixed haplotype sites in their mitochondrial genomes, ALR, FVB, and NZB. We find that in ALR and FVB strains, haplotype sites were among the most mutated sites in the mt-genome, with 6-fold and 7-fold higher average mutation frequency than non-haplotype sites, respectively **(Fig. 6B**, adjusted p-values <0.001). In NZB, which differs from B6 at 91 locations, these haplotypes sites had 122-fold higher mutation frequency than background (adjusted p-values <1×10^-53^). The overwhelming majority (75% - 100%) of these mutations are reversions to the B6 allele with the specific reversion mutation occurring significantly more than expected by chance **(Table S7)**. These results demonstrate the extreme selective pressures impressed by nuclear-mitochondrial matching to reintroduce the ancestral allele.

We next hypothesized that if somatic selection were to favor reversions, these B6 reversion alleles should increase in frequency with age. We quantified the relative change in reversion frequencies across strains and found that in ALR and FVB all unique fixed sites exhibited a significant increase in the reversion allele frequency with age across all tissues **(Fig 6C, Table S8**, BH adjusted p-value < 0.02). This suggests a strong benefit of these coding reversion substitutions in ALR and FVB strains. In contrast, in NZB we observed an overwhelming preference for reversion alleles to decrease in frequency with age, particularly in the brain **(Fig 6C, Table S8)**. One potential explanation for this decrease in frequency compared to ALR and FVB is that the many NZB fixed substitutions exhibit epistatic interactions which manifest negatively if single reversions are not accompanied by mutations at other sites, though this is challenging to test. We did identify two cases in NZB of reversions that increase in frequency with age, a synonymous substitution in the *mt-CO2* gene, and a noncoding mutation in the *D-loop*. Together, our results demonstrate that sites which contribute to mito-nuclear mismatching are prone to elevated levels of mutation with reversion alleles preferred across multiple tissues in populations. This preference for nuclear mitochondrial matching potentially drives somatic selection increasing the frequency of these alleles with age.

## Discussion

The interdependence of nuclear- and mitochondrially-encoded genes has significantly shaped patterns of diversity and constraint on these two genomes across organisms and populations. Yet, the mt-genome also exhibits exceptionally high somatic mutation rates and is thus dynamic within individuals’ lifespans. Here, we explored the relationship between nuclear-mitochondrial ancestral matching and the accumulation of somatic mutations. Our results extended upon other works that employ highly sensitive Duplex Sequencing to characterize the mutational landscape of the mt-genome ^34–37^ and generated, to our knowledge, the largest mt-genome somatic mutational catalog to date. We profiled 2.5 million mt-genomes across 4 conplastic mouse strains and the B6 wildtype, with a median of more than 74,000 mt-genomes per biological condition. Our results corroborated other studies of somatic mutation in the mt-genomes of mice, humans, and macaques, which demonstrate an age-associated increase in mutations as well as tissue-specific mutation rates. Our study design also uniquely allowed us to assess the role of mt-haplotype on mutation rates. The conplastic strains employed each exhibit distinct physiological differences which likely play a role in modulating mutations. We find cases in which mt-haplotype impacts mitochondrial mutation rate, which emphasizes that specific fixed substitutions between mitochondrial haplotypes can play a vital role in shaping mutational profiles.

Our study design allows us to compare mutational landscapes across three of the most metabolically active tissues: the brain, heart, and liver. We show that SNV mutation rates are haplotype- and tissue-specific. We also noted that while most mutations increase in frequency with age, ROS-associated mutations do not. These findings recapitulate prior studies conducted by Arbeithuber et al. 2020^35^ and Sanchez-Contreras et al. 2023^37^. Several hypotheses have been proposed to explain this phenomenon ranging from potential molecular mechanisms that recognize and remove ROS-damage through aging^37^ to the existence of mitochondrial subpopulations that dictate the replication of specific, mutation-limited mtDNA molecules^51,52^. Further work is needed to identify the ultimate mechanisms that result in such non-clocklike signatures.

Our high duplex coverage allowed us to characterize mutational hotspots throughout the mt-genome at fine-scale resolution. As expected, the *D-Loop* had a high mutation frequency compared to other regions in the genome. However, we identified three additional hotspots exhibiting either age- or haplotype-associated mutation frequencies. The first of these hotspots occurs in the *OriL,* the light strand origin of replication, which forms a DNA stem-loop structure. While the sequence of this region is highly divergent among species, the structure is conserved. Recently, Arbeithuber et al.^36^ identified the *OriL* as a hotspot of mutation in macaque liver. We were also able to capture these hotspots in a reanalysis of duplex sequencing data from several different tissues that was recently published despite their relatively lower coverage **(Fig S11 A-B, Table S9)** ^37^. Similarly to Arbeithuber et al^36^, we find mutations at the *OriL* locus to be most prominent in aged wild-type liver, however our increased sequencing depth allows us to show that this is a consistent hotspot across tissues and strains regardless of age. We identify mutations in both the stem and the loop of this structure, with mutations occurring in the same 5’ end of the stem as those identified in macaque, albeit impacting completely different sequences. We further noted that there is a well supported mechanism for this specific mutational hotspot occurring on one arm of the stem loop structure due to replication priming by an RNA which is susceptible to slippage^53^. These results highlight that the conserved structure of this stem loop is sensitizing to conserved mutational patterns across species. Similarly, peaks of mutation frequency within the *D-Loop* have been narrowed to regions associated with mtDNA replication^20,50,54^.The mutational hotspots surrounding mtDNA origins of replication have been hypothesized to serve as a compensatory mechanism against mutation by slowing mtDNA replication^36^. We identified an additional hotspot occurring in a highly structured nucleic acid, *mt-tRNA^Arg^*. Intriguingly, these mutations are only found in conplastic strains, suggesting that they are poorly tolerated in the wildtype. These mutations are expected to increase the size of the loop in the 5′ D-arm stem-loop of this tRNA. Polymorphisms that increase the size of the D-arm in *mt-tRNA^Ar^*^g^ have been linked with an increase in mtDNA copy number triggered by heightened reactive oxygen species production^55^. Specifically, the size of the loop, not the underlying sequence, was found to be important for mitochondrial function^42^. Together, these findings suggest that mutational hotspots in the mt-genome may potentially act as compensatory mechanisms to alter mtDNA replication and copy number.

While Duplex Sequencing approaches have allowed us to identify processes associated with DNA replication as the primary driver of somatic mutations in mitochondria^34–36^, the full extent of mutational processes impacting mt-genomes remains unknown. Recent studies have determined that different species exhibit varying contributions of mutational signatures in the nuclear genomes of aging intestinal crypt cells^48^. Comparing the mutational spectra across several studies that duplex-sequence the mt-genomes of humans, mice, and macaques in various tissues, we similarly find that rodents exhibit distinct mutational profiles compared to primates. Although DNA replication error is the predominant mutation type across species, rodents exhibit a surfeit of G>T mutations in contrast to primates. While studies in humans previously concluded that transitions are the most common mutation in mitochondria, we find that the abundance of the T>C marker of DNA replication error is a species-specific effect. We are not able to determine whether this finding is the result of physiological differences between these organisms, or variability in the repair pathways of these species. Of note, we find these trends hold both for somatic mutations as well as de-novo oocyte mutations in the mt-genome, suggesting that the underlying mechanisms driving these distinct profiles have potentially shaped patterns of mt-diversity across different species. Intriguingly, a very similar signature distinguishing mice and ferrets from other mammals was found using laser-capture and deep sequencing of intestinal crypts across 16 species^48^. Together, these findings thus recapitulate that distinct life history traits impact the evolution of the mt-genome across species. Compared to primates, rodents have a shorter lifespan and a substantially smaller body size. The difference in G>T mutations, which are associated with reactive oxygen species damage, and T>C mutations, a marker of DNA replication error, suggest that repair mechanisms or ROS defenses may differ between these species. Further analyses across broad ranges of species, such as those recently performed in the nuclear genome^48^, are needed to inform how mitochondrial mutational processes are specifically associated with disparate life histories.

Mitochondrial genomes exist as a population inside cells, where selection can act to shape this population at various biological scales. In oocytes, strong genetic drift induced by the mitochondrial genetic bottleneck^26^ and purifying selection^20,56^ have been shown to shape the transmission of mt-genome mutations across generations. In somatic tissues, the population of mt-genomes may be shaped at the cellular level, as a result of inter-cellular competition; at the inter-mitochondrial level, with mitochondrial turnover; or at the intra-mitochondrial level between mt-genomes harboring different variants. Previous studies focused on *de novo* mutations in mice have not identified signals of selection^35,37^. However, it has been suggested that this may be due to the low frequencies of these variants, which prevents them from having a phenotypic effect on mitochondrial function. By contrast, studies focused on higher frequency mutations in humans (greater than 0.5% frequency) have identified signals of positive selection^50^, and potential signals of negative selection^50,57^ based on differences between polymorphic and heteroplasmic nonsynonymous mutations. Furthermore, signatures of positive selection in the liver have been previously reported in both macaques^36^ and humans^50^. Complementing these studies, we find that low frequency mutations, which are likely *de novo* somatic mutations, are not under strong selection. However, mutations at intermediate frequencies, ranging from 5×10^-5^ to 1×10^-3^, do exhibit some signatures of negative selection. While these are likely not *de novo* mutations, they demonstrate a low frequency threshold for tissue-specific selection of inherited or early developmental mutations. Importantly, our results demonstrate that mitochondrial mutational spectra vary across different frequencies and mutational hotspots **(Fig. 5B)**, which can influence signatures of selection. Together, these findings suggest that purifying selection acts on mitochondrial mutations segregating at lower frequencies.

Our study also allowed us to examine a very specific form of somatic selection: the emergence and persistence of reversion mutations that re-match the mitochondrial haplotype to its nuclear constituent. Previously, Wei et al. identified the preferred maternal transmission of variants that worked to re-align mitochondrial and nuclear ancestry in humans^20^. This result showcased that selection for reversions can occur in as short as one generation, emphasizing the strong influence mito-nuclear interactions have in shaping mt-genome diversity. We find selection for reversion mutations at even shorter timescales: within an organism’s lifespan. These mutations are extremely abundant, much more than expected by chance, and in several cases increase in frequency with age. Importantly, the change in reversion allele frequencies with age was dependent on the genetic divergence between the mitochondrial and nuclear ancestries. That is, mice with the greatest genetic divergence from the B6-mitochondrial haplotype observed the opposite trend, a decrease in reversion frequency with age. Despite mito-nuclear ancestral mismatching, NZB mice did not showcase phenotypic differences compared to B6^58^. Thus, understanding to what extent mito-nuclear matching influences somatic evolution remains to be explored. Altogether, these observations highlight a hitherto unexplored impact of mito-nuclear incompatibility, namely its potential role on the somatic evolution of tissues.

While we sequence to great depth across various tissues, we are unable to characterize the impact that cell-type plays on the evolution of the mt-genome. This analysis is of particular interest in tissues such as the heart and the brain which consist of both mitotic and post-mitotic cellular populations. Given that DNA replication error is a predominant age-associated mutational signature, sequencing at the single cell level will be pivotal in understanding the role that cell proliferation has on mitochondrial somatic mutation. Additionally, our study only examined the wildtype B6 mouse and conplastic strains with nuclear B6 backgrounds.

Reciprocal conplastic strains, in which both mitochondrial haplotypes are placed in the context of both nuclear genomes, will allow us to parse apart the different roles that the nuclear and mitochondrial genomes have in driving distinct mutational processes. Lastly, our study emphasizes the importance of comparative somatic mutation profiling in order to discern how processes that shape mutation in the mt-genome differ with life history traits. Altogether, our findings explore somatic evolution in the context of an important cellular organelle and begin to discern the various scales at which evolutionary processes act to shape the population of mt-genomes.

## Methods

### Ethics Statement

Animal use and all methods were approved by the Animal Care and Use Committee (V242-7224. 122-5, Kiel, Germany). Experiments were performed in accordance with the relevant guidelines and regulations by certified personnel.

### Data collection and sample preparation

The wildtype C57BL/6J (B6) and inbred mouse strains were obtained from Jackson Laboratory and maintained at the University of Lübeck. Conplastic strains B6-mtAKR, B6-mtALR, B6-mtFVB, and B6-mtNZB were generated and bred as described in^38^ at the University of Lübeck. Briefly, the conplastic strains were developed by crossing female mice from AKR, ALR, FVB, and NZB mouse strains with male B6 mice. Female offspring were then backcrossed with male B6 mice. After the tenth generation of backcrossing mice were deemed conplastic mice with a B6 nuclear background and their respective maternal mitochondrial haplotypes. Samples were validated by checking for their defining haplotype sites **(Table S1, Fig. S12).** The brain, heart, and liver were sampled from young (2-4 months old (mo.)) and aged (15-22 mo.) mice in each strain (n = 115). B6 young liver samples and one B6 young heart sample were omitted from the data due to possible contamination **(Fig. S12)**. Whole tissue samples were flash frozen and processed for DNA isolation using the Qiagen DNAeasy Blood and Tissue kit [ID: 69504]. All mice in this study are female.

### Sequencing

Duplex sequencing libraries were prepared by TwinStrand Biosciences (Seattle, WA) as previously described^33,34^. Sequencing was performed at the University of California, Berkeley using Illumina NovaSeq generated 150 bp paired-end reads.

### Data processing

Sequencing reads were processed into Duplex Sequencing reads and mapped to the reference mouse mitochondrial genome (mm10) using a modified version of the Duplex Sequencing processing pipeline developed by Dr. Scott Kennedy’s group at the University of Washington, Seattle. The pipeline was edited to take as input bam files formed from the NovaSeq reads using bwa mem^59^ and samtools^60^, and is available on our github. Software versions used along with all processing steps including duplex consensus sequencing generation, mapping, and variant calling can be found in the github repository referenced below. Parameters used to run the Duplex Sequencing pipeline are provided in Supplemental Note 1.

### Trinucleotide Spectra

Only *de novo mutations* were used in this analysis and were identified by filtering for mutations with an alternative allele depth < 100 and a mutation frequency < 0.01. Each mutation is scored once to create a proxy mutation frequency for *de novo* mutations. The mutation fraction is the proportion of each mutation type in a given trinucleotide context divided by the total count of *de novo* mutations for a condition. The trinucleotide and mutation type featured represent mutations on either strand. For example, ACG represents both ACG and CGT, where either a C>T or a G>A mutation has occurred **(Fig. S13).**

### Mutational Signature Extraction

Observed counts for each mutation type (excluding mutations at a frequency > 1×10^-3^) were used as input for sigfit (v 2.2), which is an R package used to identify mutational signatures. Sigfit uses Bayesian probabilistic modeling to uncover mutational signatures, as explained by Gori and Baez-Ortega^47^. Specifically, we use multinomial models to extract our signatures, which is akin to the traditional non-negative matrix factorization approaches. The optimal number of signatures was determined by extracting a range of 1-7 signatures and comparing the cosine similarity **(Fig S8)** for this range (iterations = 1000, seed = 1756). The model was then refitted with two signatures, as determined by the goodness of fit test, with 10,000 iterations (seed = 1756).

### Testing for selection

The number of nonsynonymous variants per nonsynonymous sites (*hN*) and synonymous variants per synonymous sites (*hS*) were calculated as described in ^35,50^.

We quantified mutations and mutation type proportions in three different ways: (1) all mutations, without consideration of mutation frequency or mt-genome position (2) all mutations excluding the D-Loop (3) mutation counts binned by mutation frequency, excluding the D-Loop. Mutation counts were calculated as the sum of the duplex alternative allele depth across all sites in the mt-genome. Counts were aggregated across replicates in experimental conditions **(Fig. S14A)**.

For analyses where mutation counts were aggregated across frequencies, the 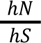 statistic was quantified for each gene across the 29 experimental conditions. To test if the observed 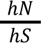 statistics were significant signals of selection, we used a multinomial distribution to simulate mutations for all experimental conditions using the observed mutation counts and proportions for each mutation type. The mutations were sampled across the mt-genome uniformly with replacement, and the simulated 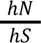 statistics were calculated for each gene. 10,000 simulations were conducted for each experimental condition. 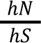 statistics that could not be calculated (*hS* = 0 or *hN* = 0) were excluded from the analysis . Mutations with a frequency > 1×10^-3^ were excluded from these analyses **(Fig. S14B)**. Empirical p-values were used to determine if an observed 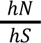 statistic was significant. The empirical p-value was calculated as the number of simulated 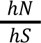 statistics with a more extreme value than the observed 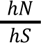 statistic divided by the total number of simulated statistics. These empirical p-values were multiplied by a factor of 2 in order to account for both tails of the distribution. 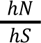 statistics with a Benjamini-Hochberg adjusted p-values < 0.01 were denoted as significant, where 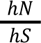 > 1 signified that a gene was under positive selection and 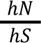 < 1 suggested a gene was under negative selection.

For analyses where mutation counts were binned by frequencies, mutations with a frequency > 1×10^-3^ were included. 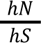 statistics were calculated for each bin across every experimental condition. Mutation counts were calculated as the sum of the duplex alternative allele depth across all sites in the mt-genome within the frequency bin. Counts were aggregated across replicates in experimental conditions. The analysis described above was performed for every bin in each experimental condition. A Benjamini-Hochberg multiple hypothesis test correction was applied for each bin to maintain consistency in the number of tests corrected for between the aggregated and binned analyses.

### Comparison of the null and observed mutational spectra for missense and synonymous mutations

The null mutational spectra for synonymous and missense mutations was calculated by quantifying the number of missense and synonymous mutations each mutation type could produce. The proportion was calculated as the count of missense or synonymous mutations resulting from a given mutation type divided by all possible missense or synonymous mutations in the mt-genome **(Fig. S15A).** For the observed mutational spectra, the proportion of each mutation type comprising the total count of missense and synonymous mutations was calculated **(Fig. S15B)**.

### Estimation of and correction for NUMT contamination

We originally mapped duplex paired end reads to a *mm10* masking the known NUMT region. We reasoned that given the 10-100-fold higher somatic mutation rate of the mitochondrial genome along with its several-hundred fold increased copy number with respect to the nuclear genome, that nuclear contaminating reads contribute minimally to our mutation frequencies. However, this low-level of contamination can potentially cause an issue when examining reversion mutations which are expected to be enriched for B6 allele in the NUMT.

To estimate and correct for this NUMT contamination, duplex paired end reads were remapped to mm10 using *bwa mem*. For this remapping, the NUMT region in chr1 (nt24611535 to nt24616184) was unmasked. The duplex read depth at junction regions, which captured sequences 10 bp upstream and 10 bp downstream of the NUMT region in chr1 and the corresponding region in the mt-genome (nt6394 to nt11042), was calculated using *samtools depth {input.in1} -b {input.in2} > {output}.* These junction regions contain sequences unique to chr1 and chrM, allowing us to estimate the % of reads mapping to ch1 as the number of reads mapping to chr1 divided by the average duplex depth of the mt-genome calculated with the chr1 NUMT masked **(Fig S16A).**

To validate this estimated contamination, we remapped the duplex paired end reads to the NZB reference genome (generated using an in-house script). The NZB mt-genome and B6 NUMT region differ by 24 SNVs. To map the duplex reads to the NUMT region we used *samtools depth {input.in1} -bq 30 {input.in2} > {output},* setting a strict mapping quality score given that reads may differ from the regions by as few as 2 positions. Six regions across the NUMT were identified as having multiple SNVs within a read (<130 bp apart), which we refer to as SNV clusters. We used these clusters to identify reads that mapped to chr1. The estimated % of NUMT contamination at each SNV cluster was calculated as the number of reads mapping to chr1 divided by the average duplex depth of the mt-genome calculated with the chr1 NUMT masked. The estimated % of contamination at the junction regions was compared to the distribution of estimated % of contamination for the SNV clusters in NZB **Fig S16B).** The maximum estimated % of contamination between the junction regions was consistently equal to the median estimated % of contamination for the SNV clusters in NZB, verifying the consistency of the estimated % of NUMT contamination.

We take as a conservative measure the maximum estimated % of contamination from the junction regions (∼0.5% contamination). The estimated chr1 read depth was then calculated as the original average mt-genome duplex depth multiplied by the estimated % of contamination. We calculated the corrected duplex mt-genome read depth in this region as the original duplex mt-genome read depth subtracted by the estimated chr1 read depth count. Likewise, the reversion allele depth is calculated as the original reversion allele count subtracted by the estimated chr1 read depth, assuming all reads from chr1 contain the B6 allele.

### Statistical analysis: Testing for significance

Statistical analyses were performed using R (4.1.2). Poisson confidence intervals for average mutation frequencies were calculated using the *qchisq* function from the *stats* package in R (v 4.1.2). Log-link regressions were performed to determine an age-associated change in mutation frequency and mt-haplotype specific mutation rates (*glm, stats* package). Associations between (1) mutation type and age and (2) reversion allele and haplotype site were determined via Fisher’s exact tests (*fisher.text*, *stats* package). To compare (1) the average mutation frequencies between the D-Loop and other regions in the mt-genome and (2) the average mutation frequencies of simulated and observed 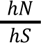 statistics on a gene-by-gene basis we used two tailed t-tests (*t.test*, *stats* package). To determine that our simulated 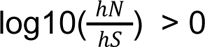 altogether and in a gene-by-gene analysis we used a one-sample t-test (*t.test*, *stats* package). To compare (1) the average 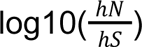 between our simulated and observed data and (2) background to haplotype site mutation frequencies for the NZB conplastic strain, we conducted Wilcoxon-Rank Sum tests (*wilcoxon.test, stats* package). Lastly, we calculate empirical p-values to determine the significance of observed 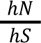 statistics; high haplotype site mutation frequencies for AKR, ALR, and FVB; and for changes in mutation frequency with age for all haplotype sites. The empirical p-values are calculated as the number of simulated 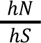 statistic (for 1) or number of background sites (for 2 and 3) with a more extreme value than our observed 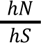 statistic (for 1), haplotype site mutation frequency (for 2), or change in mutation frequency for the haplotype site (for 3), divided by the total number of simulations (for 1) or total number of background sites (for 2 and 3). Multiple hypothesis test corrections were performed using the Benjimi-Hochberg correction (*p.adjust, stats* package), adjusted p-values refer to p-values that have undergone multiple hypothesis correction.

## Data availability

All raw data has been uploaded to sequence read archive (accession pending). Processed duplex reads are deposited in zenodo.

## Code availability

All code have been deposited in github (https://github.com/sudmantlab/conplastic_mt_profiling) with the repository archived in zenodo at (https://doi.org/10.5281/zenodo.8436686).

## Author contributions

Conceived the experimental design: PHS.

Constructed the conplastic strains and provided experimental samples: MH and SI. Sequencing design and library preparation: CV, SA, ES, GP, LW, JS. LW and GP contributed to this project while affiliated with TwinStrand Biosciences.

Processed and analyzed the data: IMS and PHS. Wrote and edited the manuscript: IMS and PHS.

